# A Highly-Efficient, Scalable Pipeline for Fixed Feature Extraction from Large-Scale High-Content Imaging Screens

**DOI:** 10.1101/2023.07.06.547985

**Authors:** Gabriel Comolet, Neeloy Bose, Jeff Winchell, Alyssa Duren-Lubanski, Tom Rusielewicz, Jordan Goldberg, Grayson Horn, Daniel Paull, Bianca Migliori

## Abstract

Leveraging artificial intelligence (AI) in image-based morphological profiling of cell populations is proving increasingly valuable for identifying diseased states and drug responses in high-content imaging (HCI) screens. When the differences between populations (such as a healthy and diseased) are completely unknown and undistinguishable by the human eye, it is crucial that HCI screens are large in scale, allowing numerous replicates for developing reliable models, as well as accounting for confounding factors such as individual (donor) and intra-experimental variation. However, as screen sizes increase, challenges arise including the lack of scalable solutions for analyzing high-dimensional datasets and processing the results in a timely manner. For this purpose, many tools have been developed to reduce images into a set of features using unbiased methods, such as embedding vectors extracted from pre-trained neural networks or autoencoders. While these methods preserve most of the predictive power contained in each image despite reducing the dimensionality significantly, they do not provide easily interpretable information. Alternatively, techniques to extract specific cellular features from data are typically slow, difficult to scale, and often produce redundant outputs, which can lead to the model learning from irrelevant data, which might distort future predictions. Here we present ScaleFEx℠, a memory efficient and scalable open-source Python pipeline that extracts biologically meaningful features from large high-content imaging datasets. It requires only modest computational resources but can also be deployed on high-powered cloud computing infrastructure. ScaleFEx℠ can be used in conjunction with AI models to cluster data and subsequently explore, identify, and rank features to provide insights into the morphological hallmarks of the phenotypic categories. We demonstrate the performance of this tool on a dataset consisting of control and drug-treated cells from a cohort of 20 donors, benchmarking it against the state-of-the-art tool, CellProfiler, and analyze the features underlying the phenotypic shift induced by chemical compounds. In addition, the tools generalizability and utility is shown in the analysis of publicly available datasets. Overall, ScaleFEx℠ constitutes a robust and compact pipeline for identifying the effects of drugs on morphological phenotypes and defining interpretable features that can be leveraged in disease profiling and drug discovery.

## 1. Introduction

High-content imaging (HCI) is a popular method to generate large, information-rich imaging datasets of cell phenotypes^1^. HCI can identify phenotypic differences between groups, such as healthy and diseased states, as well as the effects of pharmacological interventions or genetic manipulations^2^. However, unbiased analysis of HCI data requires large datasets and sufficient replication to account for confounding factors such as donor identity and experimental parameters^3,4^. As datasets grow in size, greater computational resources and more advanced analytical approaches are required^5–7^.

One approach to detect phenotypic differences is to reduce images to a set of features derived from the latent space of convolutional neural networks^5–8^ or autoencoders^9^. Such embeddings retain the predictive power of each image while ignoring noise. However, it has proven challenging to interpret these ‘black box’ deep learning models^10^. One of the first approaches to morphological characterization was to compare the distribution of measured features that could define populations of cells based on preformed hypotheses^11–13^. While this approach has been demonstrated to work well, it is inherently biased to specific features of interest and thus unable to capture phenotypes not specifically quantified. One approach to overcoming this is to use Fixed Feature Extraction (FFE)^14^, in which quantifiable features, not linked to prior knowledge, are computed across the entire dataset, resulting in a very large set of features defining the dataset beyond preconceived hypotheses. With the increase in the number of measured features, it also becomes much harder to compute a comprehensive set in a timely manner. The most popular FFE pipeline is CellProfiler^12^, an open-source tool designed for high-throughput image analysis, allowing the extraction of quantitative information from microscopy images of cells, tissue, and other biological samples. As the first, large, open source tool of its kind, it features an intuitive graphical user interface that enables the creation of custom pipelines with cascaded calculations performed on a set of multichannel images, empowering those without programming expertise and significantly broadening the field. The creators have provided the community with pre-made pipelines, such as the widely used Cell Painting assay^15–17^, within a highly customizable tool. Using a set of dyes to label specific cellular components (Nucleus (DNA), Nucleoli and Cytoplasmic RNA, Endoplasmic Reticulum, Actin cytoskeleton, Golgi and Plasma membrane, and mitochondria), Cell Painting allows scientists to image parts of the cell across 5 fluorescence channels. Unfortunately, scaling this tool to datasets hundreds of thousands of images (or larger) in size poses computational and cost challenges. The creators of CellProfiler have addressed some of those issues by providing a containerized cloud implementation that performs distributed parallel computing over multiple virtual machines and optimizes the processes with respect to cost or time^18^. While this approach offers significant flexibility, it also has some drawbacks and can be costly. For instance, running distributed CellProfiler requires users to edit more than 5 different configuration files, each requiring extensive knowledge of the functioning of the modules, as well as experience with image analysis and cloud services. Furthermore, the authors acknowledge that performance decreases when processing more than 10,000 fields of view^19^, posing significant limitations on scaling to larger screens and potentially increasing the already considerable costs^20^.

Beyond the technical issues, the resulting output (feature vector) contains thousands of features (over 3,000 for a Cell Painting assay), many of which are redundant and highly correlated. For example, in Tegtmeier et al. ^3^ it was shown that 93% of the traits had a Pearson coefficient >0.9. This can lead to computational limitations, feature ranking difficulties, and the risk of overfitting, especially when dealing with datasets containing hundreds of thousands of cells. To address the challenges of feature extraction in large HCI datasets, we developed ScaleFEx℠, an efficient Python pipeline for extracting various FFE cell features at the single-cell level. ScaleFEx℠ is a user-friendly, scalable tool requiring minimal computational and coding expertise, and supports cloud deployment. It calculates features such as cell shape, cell size, pixel intensity, texture, granularity, zernike moments, correlations between fluorescence channels. Additionally, ScaleFEx℠ can measure features that are specifically relevant to mitochondria and RNA by adjusting a coded flag. These features are crucial for studying cellular dysfunctions linked to a variety of diseases. ScaleFEx℠ produces interpretable, lightweight representations of single-cell images, aiding in the separation and characterization of cell populations using AI. We demonstrate ScaleFEx℠’s utility by identifying phenotypic shifts in drug-treated cells and validating these shifts through feature analysis and image correlation. Additionally, we showcase its generalizability by analyzing a public dataset (RxRx2)^21^ to compare morphological effects of small molecules. ScaleFEx℠ offers a novel, computation-friendly approach to extracting interpretable features from HCI datasets, aiding scientists in understanding biological changes related to disease and drug treatment.

## 2. Results

The primary goal of ScaleFEx℠ is to efficiently compute interpretable features from images of single cells stained for multiple cellular structures to characterize their morphology in large scale experiments (Figure 1), while also being easy to use and adaptable to different immunohistochemistry panels. We developed the pipeline with the Cell Painting assay in mind, but it can be used on any staining panel, as long as a nuclear stain is provided. The selection of features was guided by the need of interpretability and reduced output size. Our aim is to capture a comprehensive set of features that preserves essential morphological information while mitigating the challenges posed by high-dimensional data (a full description of all the features is in Supplementary Table 1). Unlike traditional pipelines, which often suffer from excessive redundancy, ScaleFEx℠ emphasizes efficiency by computing features on a per-channel basis. For example, the CellProfiler CellPainting pipeline first divides the cell into three main compartments—Nuclei, Cytoplasm, and Cell— based on the Hoechst, RNA, and Hoechst + RNA channels, respectively. It then loops over each channel to compute the same features for each compartment. In contrast, our approach computes each feature based on the channel itself, which results in approximately half the total number of features. This targeted feature computation not only reduces computational overhead but also enhances processing speed and minimizes data redundancy, making our pipeline significantly more efficient and scalable.

**Figure 1.**
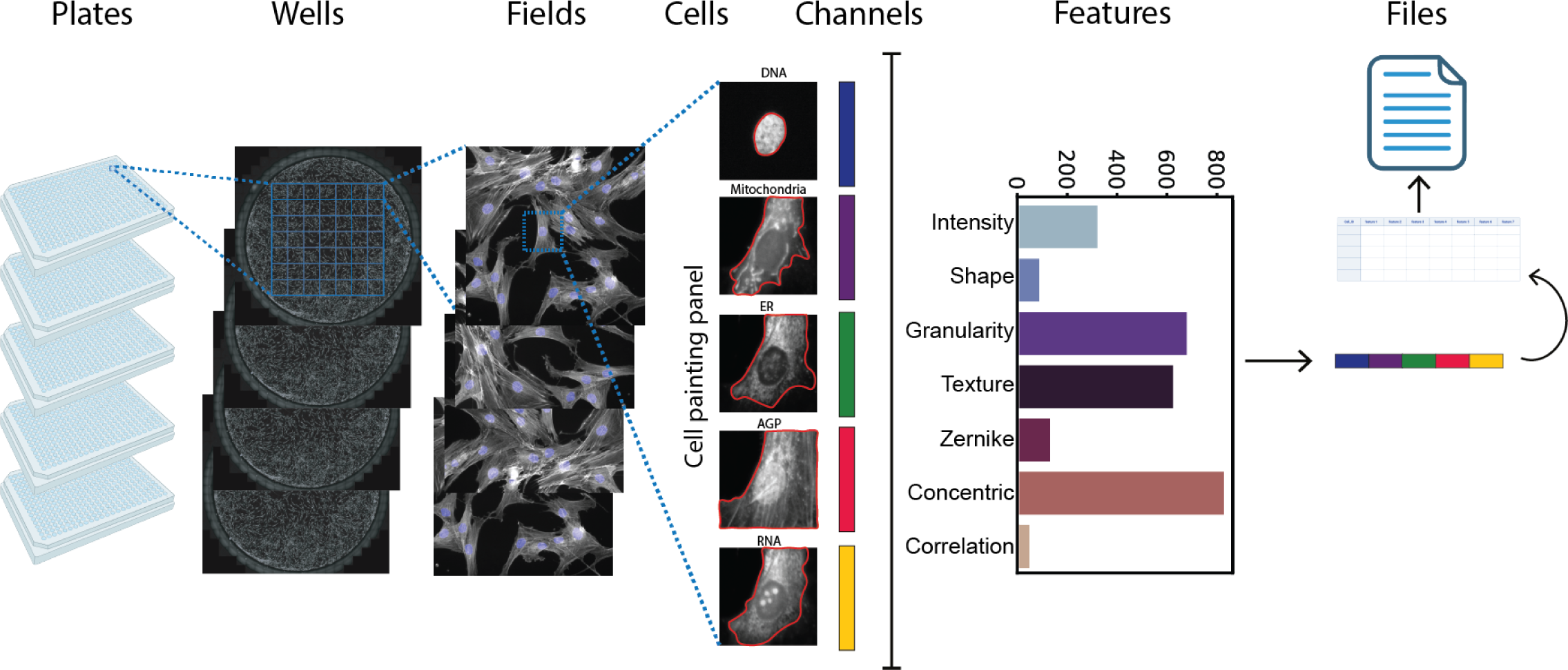
Overview of ScaleFEx℠’s pipeline on a CellPainting assay. For each experiment, ScaleFEx℠ queries the plate’s images and starts a serialized computation (parallelized over multiple machines per plate in the AWS version). Next, the wells are distributed and computed in parallel according to the number of workers specified by the user. Within each worker, after locating the centroids of the nuclei, each channel image is loaded into memory and cropped around the centroid. For each image a segmentation mask is computed, and starting from there all of the subsequent channel based features. Once the full feature vector is computed (containing measurements related to Shape, Texture, Zernike moments, Granularity, Intensity, Concentric rings, Colocalization, Mitochondria and RNA), the vector is appended to the experiment’s csv file, to avoid needing to store the entire dataset into the Random-access memory.

ScaleFEx℠’s architecture was designed with computational power limitations in mind, ensuring stability and resource optimization. The pipeline, written in Python, loops over plates, wells, fields, single cells, and channels in a hierarchical way (Figure 1, Supplementary Figure 1a), while keeping the memory requirements low. Leveraging parallel processing across a flexible number of CPUs, this method enables the independent and simultaneous computation of multiple wells, increasing performance and optimizing it for any type of resource available. By leveraging cloud services we are able to demonstrate a beginner-friendly and cost effective way to compute the ScaleFEx℠ pipeline on virtual machines. This implementation offers the option to scale to a virtually infinite number of plates by provisioning multiple machines to compute plates or subsets of plates in parallel. The parallelization process is set up to automatically connect our software with cloud-based resources, minimizing user intervention and AWS specific knowledge requirements. ScaleFEx℠ maximizes the available resources by allowing the user to adjust the number of workers used for simultaneous computation, which reduces overhead and maximizes resources, making this approach also cheaper when deployed on AWS (Table 1). (See Methods section, code-repository and associated wiki for the documentation and implementation strategy of the pipeline to deploy ScaleFEx℠ on a cloud-based platform for users that might have extremely large datasets or insufficient computational power).

**Table 1:** Comparative Analysis of Feature Extraction Tools. Comparison between ScaleFEx℠, CellProfiler, and an embeddings approach across various metrics including computation time and cost on AWS, number of features, and data correlation characteristics for a single plate. It details the performance of these tools in binary drug classification as measured by the average Area Under the Curve (AUC).

|  | ScaleFEx<sup>SM</sup> | CellProfiler | Embeddings |
| --- | --- | --- | --- |
| Computation time for 1 plate (AWS) | 35' | 1h 50' | NA |
| Computation cost for 1 plate (AWS) | ~3\$ | ~40\$ | NA |
| AWS machine infrastructure for 1 plate | 6* C5.12xlarge (288 VCPU) | 200* C5.xlarge (800 VCPU) | NA |
| Output size | 16 GB CSV/<br>8GB Parquet | ~ 100 GB CSV | 2.6 GB CSV/1.6GB Parquet |
| Aggregation step | None | 5h | None |
| File download cost | \$0.8 | ~ \$10 | NA |
| Total number of features | 1861 | 3578 | 320 |
| Uncorrelated features | 325 | 413 | 320 |
| Number of 0 variance features | 0 | 36 | 0 |
| Total used features | 325 | 377 | 320 |
| Average AUC for binary drug classification | 0.91 | 0.88 | 0.92 |

The ScaleFEx℠ code outputs a Comma Separated Values (CSV) file (of ∼2.5 GB per plate for a typical experiment), where each row contains measurements from each cell (one row per cell). The cell measurement vectors are appended to the CSV file at each iteration, to avoid the need to retain the entire table in memory, allowing us to release it incrementally and enhance computational efficiency.

With very limited hardware requirements, ScaleFEx℠ can be deployed on any computer. Even though performance increases with the use of multiple CPUs, ScaleFEx℠’s well-parallelization and lightweight implementation significantly reduces the run time compared to tools such as CellProfiler when run on similar devices.

ScaleFEx℠ does not incorporate a graphical user interface (GUI). This decision minimizes overhead and reduces the consumption of background resources, ensuring that our tool remains lightweight and fast. Instead of a GUI, users interact with the tool through a YAML configuration file. This approach is tailored to low to mid-level Python users who have a grasp of basic programming concepts but may not be experts in software development. The initial setup requires the user to specify approximately ten run-specific parameters in the YAML file, and a few that can be left as default, designed to be straightforward. The parameters are clearly defined and documented, making it easy for users to understand what information is needed and why it’s important. After the initial setup, for subsequent runs, users typically only need to update the folder locations if the experimental settings remain unchanged. This simplifies repeated use and reduces the potential for errors, as most of the configuration is already completed and verified. By using a YAML file for configuration, we provide a user-friendly way to input necessary parameters without navigating complex software interfaces. YAML is chosen for its human-readable format and its ability to clearly represent hierarchical and scalar data, making it particularly suitable for configuring software settings. We also incorporated some visualization options to help verify that the parameters are correctly set before launching the tool over a very large dataset.

The AWS implementation is also designed to minimize the user interaction with the sometimes complex AWS infrastructure. The initial steps involve setting user permissions and creating a secure environment through AWS’s IAM and CloudFormation services. Each run is easily managed by executing predefined templates that automatically handle the setup and teardown of necessary resources. This straightforward process ensures that even users new to cloud computing can efficiently manage and scale their computations without unnecessary complexity.

### 2.1 Benchmarking

To benchmark ScaleFEx℠ we used a dataset consisting of fibroblasts from 20 donors treated with three compounds at different concentrations, stained using the Cell Painting panel and imaged at high resolution, resulting in a dataset of 294,000 images (2.5 terabytes). Every step of the plating of the cells into the 5 384-well plates was conducted using automation as part of the NYSCF Global Stem Cell Array®^22^, (as described in the Methods section and Supplementary Figure 1b). We extracted ScaleFEx℠ and CellProfiler features and calculated image embeddings (vectors encoding abstract image features calculated using deep neural networks sas described in Schiff et al.^7^). Running feature extractions algorithms such as CellProfiler over a large number of images poses great challenges, especially since GPU computation (typically used to accelerate image-related analyses) is mainly convolution based and is not optimal for complex logic operations and measurements with sequential dependency between the features extracted or accumulative computations ^23–25^. To evaluate the performance of ScaleFEx℠ and CellProfiler, we carried out a series of benchmark tests (summarized in Table 1). We first processed a subset of our data (8 wells, 1960 images) on a desktop machine. CellProfiler completed the task in 4 hours, while ScaleFEx℠ took only 1 hour. The efficiency of ScaleFEx℠ was significantly enhanced by its ability to process multiple wells concurrently and its reduced feature computation load, dramatically cutting down overall computation time.

We next compared the cloud based performance of ScaleFEx℠, deploying it to AWS and compared this to the distributed cell profiler^19,20^ across the entire dataset. Distributed CellProfiler is designed to parallelize AWS machines over images, hence the machines recommended to be deployed are small and cheap. With this approach, a large number of machines need to be deployed, but it is easy to run into limits of availability. The authors also recommend not to deploy more than 200 machines, and caution that the process slows down with datasets comprising over 10,000 fields^20^. In addition, each machine requires initialization, package installation and processing of the first steps. ScaleFEx℠’s AWS implementation leverages parallel processing across both individual machines and their CPUs, optimizing configurations with slightly larger machines than those used by CellProfiler. This approach not only shortens deployment times but also reduces costs significantly. Additional cost efficiency is gained by utilizing Spot Instances, where users can bid on machine time at their desired price, thereby controlling expenses and avoiding unexpected charges. When comparing the cost and time of analyzing the entire dataset, we observed a large difference in performance with a ten fold reduction of costs and 4 times faster computation when using our pipeline (Table 1). CellProfiler’s output came in various different files per plate that had to be merged based on the information of plate, field and cell ID, a step that can require a long time to process and implement (∼5h for the dataset we described in this paper) (Table 1). The final process leads to a vector of 3578 numbers per cell, which, multiplied by the large number of cells in the entire dataset, was challenging to load into memory, merge, and process. For this reason, we averaged the features at well-level as the authors of CellProfiler do on most of their analyses^2^ before merging the files, as this operation was saturating available memory. ScaleFEx℠, in contrast, creates a feature vector of 1861 total features for the same channels and cells. This dataset can be loaded in under a minute, allowing one to perform multiple operations on the cell-level features, since no merging is required and the overall occupied memory was around half that of CellProfiler.

To compare redundancy between CellProfiler and ScaleFEx℠, we performed a correlation analysis on the same dataset and counted the remaining features after removing all the features that had a Pearsons’s correlation score above 0.9 (this threshold was taken from the value chosen in Tegtmeyer et al. 2024^3^). CellProfiler had a slightly higher percentage of correlated features compared to ScaleFEx℠ (Table 1), and 36 of features that had zero variance compared to 0 features in ScaleFEx℠. Both methods had a high number of correlated features, which is expected in this type of measurement.

To further compare CellProfiler’s output to ScaleFEx℠’s, we used a Logistic Regression (LR) model to assess the predictive power of each on the same task. We also compared the performance of feature-extraction-based (both ScaleFEx and CellProfile) with deep learning-based dimensionality reduction approaches by evaluating the deep embeddings in the same manner. We built a binary classifier trained to predict if a well was treated with a compound or not, for each compound and condition of the dataset (for details see the Methods section). First, we corrected for different known confounders on our data, both individually and in combination. Then, we tested three normalization methods on this adjusted data: applying a whitening transformation^26^ to decorrelate and standardize variance, z-scoring to achieve a zero mean and standard deviation 1, and normalizing features between 0 and 1. The results showed that the 3 models have very similar predictive power for the task in all the combinations of normalization and confounder removal, with ScaleFEx℠ performing slightly better in most combinations than CellProfiler and the deep embeddings being slightly better overall (Table 1, Supplementary Figure 2), demonstrating that even though ScaleFEx℠ has less features than Cell Profiler to begin with, it still provides similar performance. The best overall correction method was the normalization between 1 and 0, combined with the confounder removal of plates, rows and donor ID, so subsequent analysis was performed using this method.

### 2.2 Validation of ScaleFEx**℠** on a novel compound-treated fibroblast dataset

Following the deployment of ScaleFEx℠, we performed a detailed validation of the method by analyzing its ability to extract and highlight meaningful features from a dataset of skin fibroblasts treated with three drugs (CP21R7, Pemigatinib, and Y-39983-HCl) at two concentrations (0.2 and 1.0 μM). The objective was to validate ScaleFEx℠’s efficacy in detecting nuanced changes in cellular morphology and function, thereby accurately characterizing drug-induced phenotypic shifts. Following the steps depicted in Figure 2a, we first evaluated the presence of confounding factors that might have masked or influenced the results as previously described^3,7,27^. We iteratively removed the contribution of each confounder both singularly and in combination and evaluated the prediction score using the LR model and Uniform Manifold Approximation and Projection (UMAP)^28^. We next evaluated the accuracy of a LR model built to identify each drug compared to control (DMSO) for each of the known confounders of the experiment (plates, wells, rows, columns, donor), on well level averages. The results show that the model’s area under the curve (AUC) is the highest when correcting for plates, columns, rows, and donors (Figure 2b). This is confirmed also by the UMAP clustering (Figure 2c). The correction was performed by removing the mean of each confounder from each feature column. This correction was very effective in mitigating the confounding effects due to the cells being in different plates, different rows, and from different donors, as shown in the UMAP clustering of the wells without correction (Supplementary Figure 2). Although removing column or well confounders led to lower predictions (Supplementary Table 2), this was likely due to the layouts being similar, resulting in the removal of part of the signal with the confounder. As for the removal of the donor confounder, since primary fibroblasts retain a very strong donor signature^7^, we preferred to mitigate the individual contribution at this time. We decided to focus exclusively on the effects of the drugs to ensure that the findings are robust and not skewed by the characteristics of specific donors.

**Figure 2.**
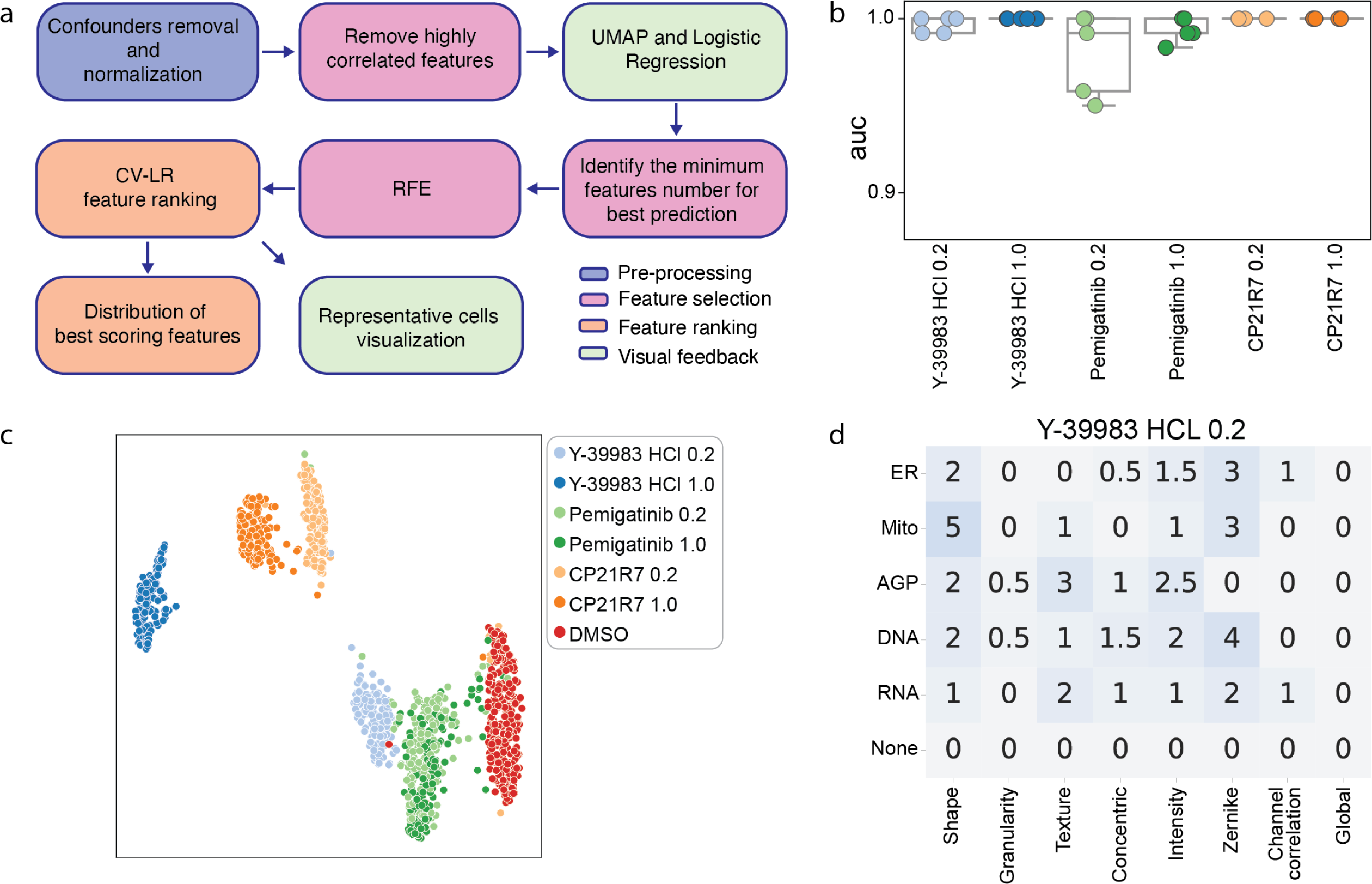
Analysis and Feature Extraction Overview. a) After a step of removal of confounders by subtraction of the mean of the confounders (plate, donor and row effects) and consequent normalization of the measurements to values between 0 and 1, the correlated features are removed at single cell level. The drug signal is evaluated by clustering the well averages of the values using the UMAP algorithm. The minimum number of features necessary for optimal prediction accuracy are identified using a LR model, and, using RFE, unimportant features are iteratively removed, leaving behind the most impactful features contributing to prediction scores. Finally, representative images of the features of interest are extracted and compared to their ranking. b) Distribution of Binary predictions of drug VS control using a Linear Regression model and 5 CV folds, where in each fold the training is performed on all the plates but one held out for testing. The model is trained to classify wells treated with each drug iteratively and DMSO controls. The values were corrected for plate, donor, column and row effects, and normalized between 0 and 1. Boxes represent median, upper and lower quartiles, the whiskers represent the standard deviation, and the diamonds the outliers. c) UMAP of the ScaleFEx℠ vector color-coded by drug and concentration. d) Example of feature summarization, by channel and category for cells treated with Y-3998-HCl at 0.2 μM. In the heatmap, the darker the color, the higher the number of features. The number states the actual number of feature per class and channel. The half numbers are for those features with double class or double channel.

The final data we used for feature analysis was confounder-corrected, normalized between 0 and 1 as previously described, and averaged at well level to reduce noise and highlight the prevalent effects in most of the population of cells^7^. We next used a Recursive Feature Elimination algorithm to eliminate the least important features for each drug based on a LR binary model built to classify each drug vs control. The most-correlating features were removed to reduce redundancy and prioritized the most important and distinct features contributing to the drug shift, while avoiding over-emphasis on similar features. The resulting features are summarized in heatmaps (Figure 2d, Supplementary Figure 3, 4,Supplementary Table 3).

To explore the phenotypes further, we analyzed the highest ranking features for each drug and examined their value distributions (Figure 3, Supplementary Figure 3, 4) to assess significant differences using a Mann-Whitney t-test. From UMAP plots and linear regression prediction accuracy, it is clear that the set of measurements is sufficient to easily separate and cluster cells treated with different compounds at different concentrations.

**Figure 3.**
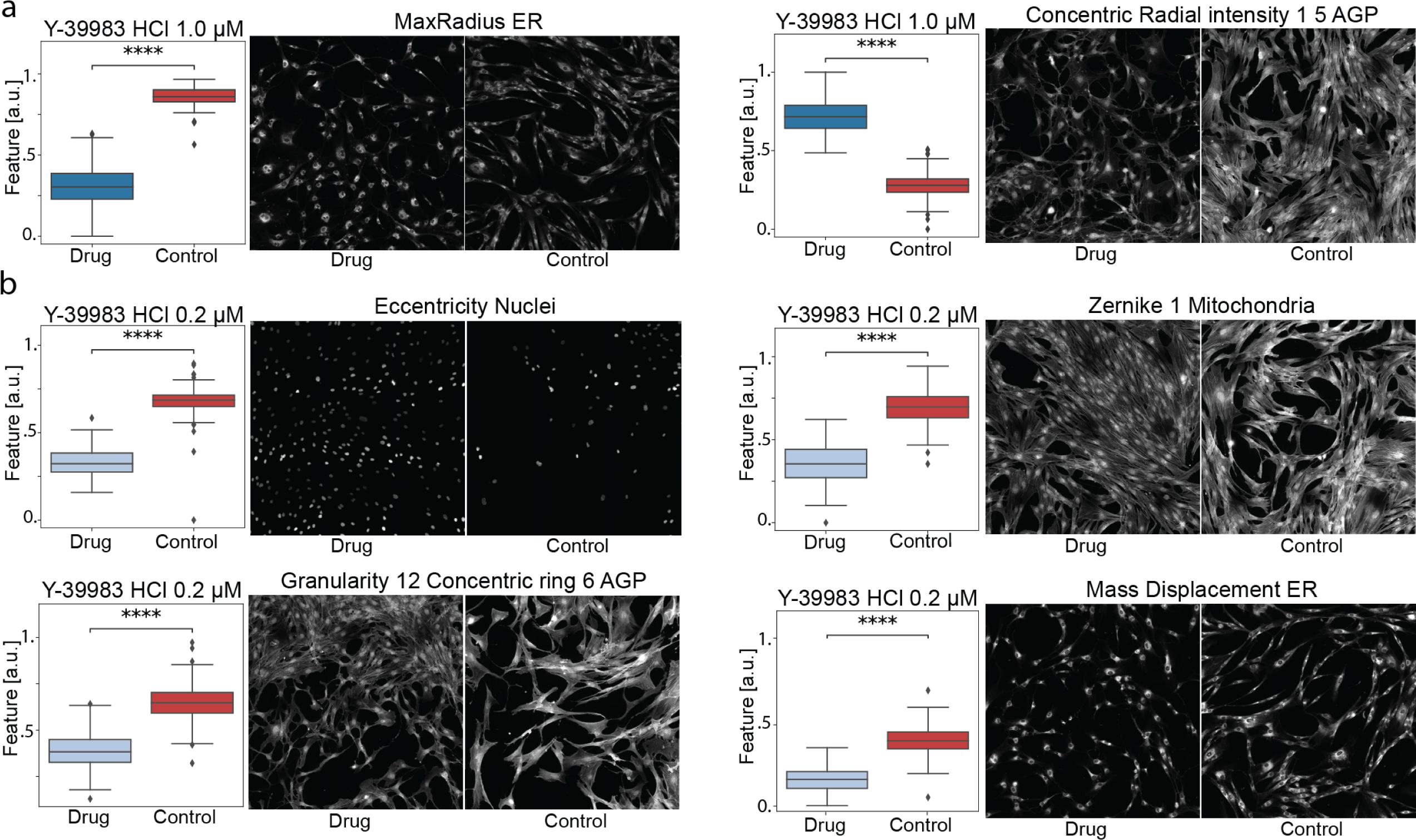
– Value distribution and image visualization of the most important features to correctly classify Y-39983 HCl. a) Features highlighted at 1.0 μM and (b) 0.2 μM. The pictures depict the wells with the closest value to the mean of the feature. In interest of space, we only visualize 9 out of the 49 total tiles that make a well. The feature distribution values are all normalized between 0 and 1. Boxes represent median, upper and lower quartiles, the whiskers represent the standard deviation, and the diamonds the outliers. **** = p < 0.0001.

**Figure 4.**
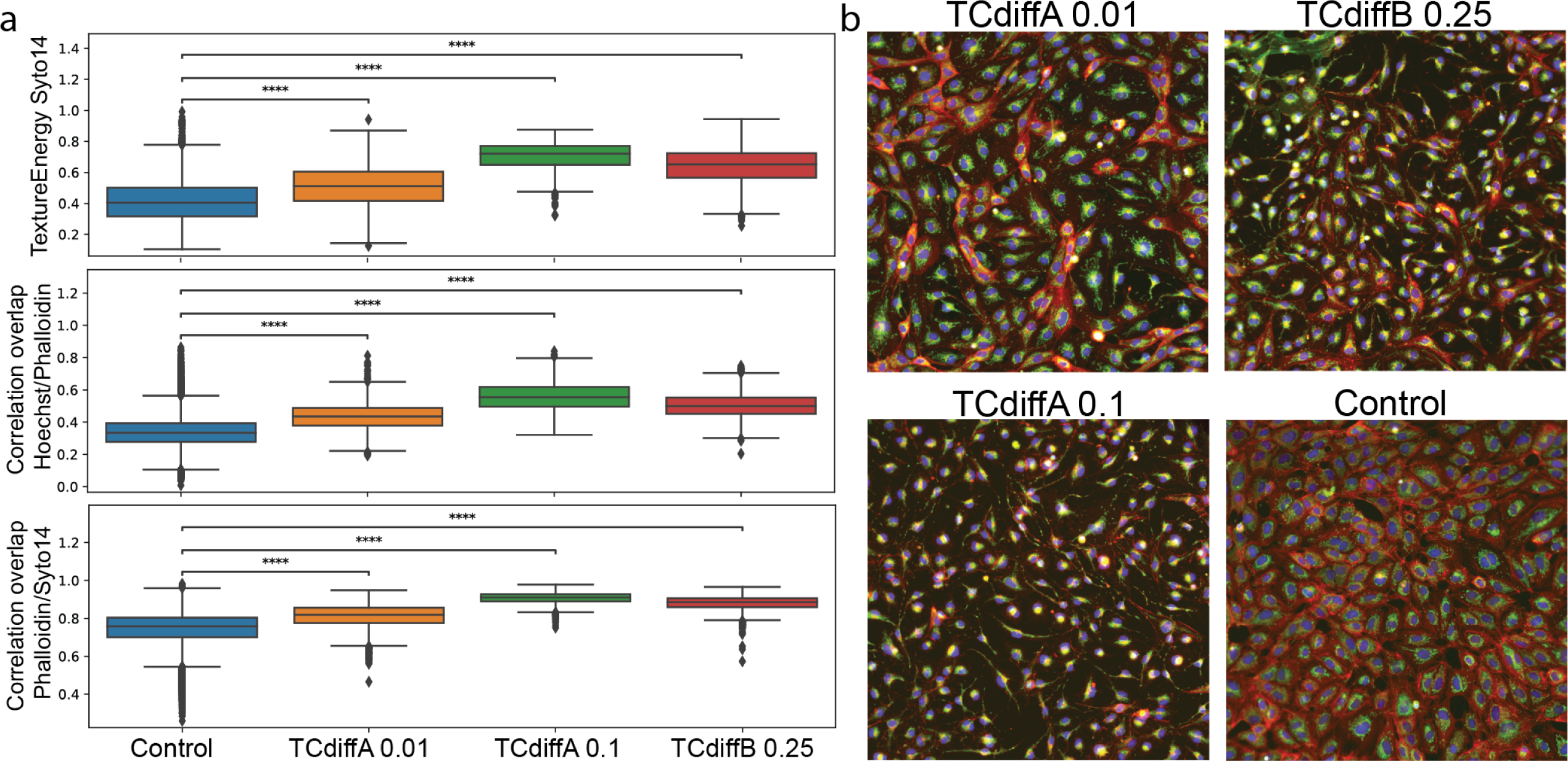
Data from one of the groups that were created using ScaleFEx℠ and similarity analysis. a) Boxplots show the most significant features in the task of separating each drug from the control and common across the 3 groups. b) Most representative tiles of the drugs that were grouped together, chosen by best separation from the control

As expected, the compounds that caused the largest shifts required fewer features to correctly classify across most wells (Figure 3a). In contrast, more subtle differences required more features (Figure 3b, Supplementary Figure, 4). This does not mean that the shift affected only those features, but that the distribution of the features is so different from the control that these features can separate the two classes.

The drug Y-39983 significantly altered cell morphology, reducing the size and altering the shape of the cell body, as indicated by features like max radius ER and concentric radial intensity in the AGP and ER channels (Figure 3a). At higher concentrations (1.0 μM), fewer features were sufficient to distinguish treated from control cells, highlighting changes such as more compact and irregular cell shapes. These findings were consistent with Y-39983’s known biological action of disrupting actin filament organization, which impacts cell motility and the cellular architecture^29^. At a lower concentration (0.2 μM), morphological changes were still detected Figure 3b) and although more subtle, were still significant, including decreased eccentricity of nuclei and altered granularity around the cell periphery, indicating changes in actin filaments and cellular texture.

Pemigatinib, a protein kinase inhibitor, caused a notable shift towards rounder cell shapes, which, amongst other metrics, caused a decrease in the form factor value in the membrane channel (Supplementary Figure 3, Supplementary Table 3). As a known inhibitor of fibroblast growth factor receptor 2 (FGFR2)^30^, Pemigatinib was also found to alter the cytoskeletal structure, causing a rounder appearance in treated cells. In addition, several mitochondrial texture features were found to be altered, suggesting an increase in mitochondrial fragmentation, typically associated with impaired cell proliferation. Both tested concentrations demonstrated similar effects, with UMAP analysis (Figure 2c) showing overlapping clustering of features and consistent, significant deviations from the control (Supplementary Figure 3).

The GSK-3β inhibitor, CP21R7, is a GSK inhibitor, a type of compound that has been shown to cause widespread disruption of cellular function including migration and cytoskeletal changes^31^. We identified changes in mitochondrial volume and organization at both tested concentrations (Supplementary Figure 4, Supplementary Table 3), confirming earlier findings that CP21R7 can alter cellular metabolism^36^. A. In addition, we found an increase in the overall size of cells stained with AGP dye, making them appear larger than other internal compartments such as the ER or mitochondria when compared to controls (Supplementary Figure 4).

### 2.3 Validation of ScaleFEx℠ using a publicly available dataset of cellular responses to immune perturbations (RxRx2)

To confirm the generalizability of ScaleFEx℠ we applied the computation pipeline on 6 plates of the Recursion dataset RxRx2^21^. This dataset encompasses 6-channel, Cell-Painted images of Human Umbilical Vein Endothelial Cells (HUVEC) from a single cell line seeded in a 1536-well plate and treated with various small molecules. The total number of cells is over 2 million, treated with 434 unique molecules at 3 different concentrations each. The analysis aims to identify drugs that induce a morphological shift compared to untreated cells, and then group these drugs by similar effects observed on the cells. We first calculated the cosine similarity for each well to quantify the similarity between cells’ multidimensional feature sets, helping to establish a baseline of cellular similarity in control wells, providing a reference point.

We next identified drugs of interest based on their lower average cosine similarity compared to the control mean, as this would be indicative of having caused changes in cell morphology or structure. We ensured that the variability within wells treated with the same drug mirrored that observed in the control wells as this would indicate that the drug’s effects are uniformly distributed across cells within a well, thus enhancing the reliability of our findings and reducing false positives due to random variation. In addition, we excluded conditions with fewer than three repeats as our analysis only utilized a subset of the dataset (Supplementary Table 4). This exclusion left us with a total of 1004 viable drugs for analysis. The entire process resulted in approximately 15 different combinations of drugs and concentrations, demonstrating significant and consistent cellular changes compared to the untreated state.

We focused on identifying drugs with similar effects on cell morphology by assessing the cosine similarity values across all remaining wells of each of the drugs. This analysis allowed us to group the remaining drugs and concentrations into 6 distinct clusters based on their similarity (Supplementary Table 4). We next explored the specific features that are more relevant for LR binary classifiers, which were built to predict drug versus control outcomes (Supplementary Table 4), and assessed whether these features were consistent across the groups. In Figure 5 we show representative images from one of the groups and some of the overlapping features that scored high for the LR model. The consistent trends observed in these features indicate that the small molecules induce similar effects on the cells, albeit with varying intensities. This observation is consistent with the known targets of TCdiffA and TCdiffB, both toxins targeting similar cellular pathways. The other groups also show similarities in the top-scoring features and representative images. Group 1 (Supplementary Figure 5) indicates a cytotoxic effect leading to cell death, while groups 2 and 3 show differences in intensities and channel overlap. To further validate these findings, we compared 2 drugs that had distinct effects identified by the algorithm and plotted their top features (Supplementary Figure 5, group 4). The standard deviation of ER intensity appears to play a role in both drugs, but their effects, as indicated by the distribution, are opposite. WGA intensity was impacted only for the Diphtheria toxin (DTx), whereas only Nigericin induced a shift towards a more elongated cell shape. Overall, this analysis was able to decipher specific features that differentiate drug effects from controls and highlight meaningful associations between the effects of different drugs on cells.

## 3. Discussion

The development of ScaleFEx℠ was driven by the need to address scalability issues common in high-throughput, automated laboratories that handle multiple cell types and assays. As the demand for high-throughput analysis grows^32–35^, laboratories require tools that can be easily adapted and deployed across diverse datasets without extensive manual configuration. ScaleFEx℠ meets this demand by offering a solution that is generalizable across any cell type and imaging method, easy to deploy both on local machines and on AWS cloud infrastructures, and scalable to adapt to different resource types.

ScaleFEx℠’s design minimizes the number of manual inputs required, allowing users to easily adjust critical parameters. This flexibility, combined with its general applicability to any cell type or imaging method, underscores its utility in a rapidly evolving research landscape where adaptability and efficiency are paramount. To our knowledge, ScaleFEx℠ processing efficiency and speed are unmet compared to tools that quantify a similar amount of measured features, making it suitable for large-scale datasets where fast feature extraction is critical.

ScaleFEx℠ is designed under the assumption that users have basic to intermediate Python skills and does not include a graphical user interface. The absence of a GUI, while beneficial for reducing complexity and improving performance, may limit its accessibility to researchers less familiar with scripting and future development of the tool may ultimately see the release of GUI. However, paramount to this will be ensuring that the performance levels gained in ScaleFEx℠ are not diminished.

ScaleFEx℠’s modular design allows for easy updates and integration of new features or algorithms, which is suitable for advanced users seeking customization within a structured framework. However, unlike CellProfiler, which allows users to tweak almost every aspect of the feature extraction process, ScaleFEx℠ does not offer the same level of detailed customization. This could be a limitation for researchers who need highly specialized analysis protocols but prefer not to engage extensively in code writing. As ScaleFEx℠ is adaptable and prepared for future enhancements due to its modular architecture, flexibility can be increased with future updates.

Currently, the majority of HCI screens extract features either using Deep Learning (embeddings or autoencoders)^1,7,21,37^ sacrificing interpretability, or selecting a small number of features to compute^38^, as an extensive representation of measurements would be extremely cumbersome or lengthy. Through this work, we aim to enhance the interpretability of HCI-based phenotyping by providing a method that not only facilitates a deeper biological understanding and explanation of subtle differences, often requiring extensive experiments to elucidate, but also ensures that it can be deployed efficiently and effortlessly.

Future developments of our platform will be focused on incorporating advanced machine learning algorithms to improve cell identification and channel segmentation. Currently, we have intentionally simplified these tasks to avoid the complexities and slowdowns associated with deep neural networks, which can also hinder parallelization capabilities and increase costs on cloud servers. Users who have already computed coordinates with centroids can seamlessly integrate these into our algorithm. This flexibility allows users to employ their preferred cell segmentation tools by either modifying the Nuclei segmentation module or by specifying the location of the file that contains the coordinates.

## 4. Conclusion

In this study, we describe the development of ScaleFEx℠, a Python pipeline that extracts fixed features in single cells, suitable for analyzing large high-content imaging datasets with minimal computational requirements. We demonstrate the utility of ScaleFEx℠ for distinguishing different drug-treated cell populations with high accuracy, and identifying salient, interpretable features that contribute to this classification process. We successfully confirmed the impact of the drugs on cells by visually comparing selected wells with control wells that best exemplified their respective medians. This approach facilitated identifying the direct link between drugs and their effects, thereby providing a transparent and reliable alternative to many AI-based dimensionality reduction tools that operate in a black-box style. Overall, this study highlights the potential of ScaleFEx℠ as a tool for identifying and characterizing drug effects on cellular populations, paving the way for the discovery of new therapeutics and disease treatments.

## 5. Methods

### 5.1 Cell collection and expansion

Cell lines used in this study were obtained through the New York Stem Cell Foundation Repository. All fibroblast cell lines derived from biopsies had been previously collected under IRB approved protocols, as previously described^22^. Cell lines were expanded and frozen into working aliquots using the NYSCF Global Stem Cell Array®, a fully automated cell culture platform, which consists of liquid handling systems (Hamilton STAR) that are integrated with (amongst other devices) automated incubators, Celigo cell imagers (PerkinElmer), VSpin centrifuges (Agilent), and Matrix tube decappers (Hamilton Storage Technologies)^7,22^. In brief, frozen master stock tubes were thawed in a 37°C water bath for 1 minute. After pelleting and resuspension in FEM, a manual count was taken, and cells were seeded into either one well of a 6 well plate, or one well of a 12 well plate in Fibroblast Expansion Media (FEM). Cells plated into 6 well plates immediately entered automated expansion workflows, while those remaining in 12 well plates were allowed to grow to 70-100% confluence before being passaged into 6 well plates and entering automated expansion workflows. Cells were fed every 2-3 days using automated scheduling and were passaged every 7 days at targeted densities across as many wells as the software calculated to be possible. Automated methods dissociated cells using TrypLE Select (12604013, ThermoFisher) before being centrifuged into pellets. The pellets were suspended and pulled into a total of 2 mL media, with two 25 µL aliquots from each sample used in automated cell counting. Based on the total live cell count, cells were frozen, using the cryoprotectant Synthafreeze (A1254201, ThermoFisher), into cryovials at 50,000 cells per vial or replated into 6 well plates for further expansion. DNA from each cell line was obtained during the process and used to confirm cell line identity using the SNPTrace™ Assay (100-6280, Standard BioTools). Cells were additionally screened for their sterility and the absence of mycoplasma during this process (LT07-703, Lonza).

### 5.2 Automated screening

Novel automated procedures were developed to run a drug screen leveraging the automated platform of the NYSCF Global Stem Cell Array®. 20 unique donor lines were selected, and a single vial of each line was automatically thawed into one well per line of a 12-well plate (Fisher Scientific, 07-200-91) for post-thaw recovery, using automated methods as previously described. Cells were fed on Days 2 and 5, and by Day 7, when the cell lines had reached over 90% confluence, the plates were passaged. At this point, the liquid handler aspirated the spent media from the seeded wells, washed them with Phosphate Buffered Saline (PBS) (Gibco™, 10010072), and incubated them with TrypLE. Cells were incubated for 25 minutes at 37°C, and then brought back to the deck for neutralization with FEM. The cell suspensions were then pipetted into an intermediate block (Corning® 3958) and centrifuged on the Agilent VSpin.

Post-centrifugation, the supernatant was aspirated, and the cell pellets were resuspended in FEM. A 10 μl aliquot of cell suspension was incubated with Hoechst (H3570, ThermoFisher) and Propidium Iodide (P3566, ThermoFisher) in a Perkin Elmer ViewPlate (502105844, ThermoFisher) before being counted on a Celigo automated imager. The counts were fed back into the automated method, allowing the determination of the required volume of cell suspension needed to seed the wells of the destination plate; a targeted density of 500 cells per well was used in this study with cells seeded in a total volume of 50 µL. The method was provided with a pre-designed, simplified, layout, and the liquid handler was able to stamp the desired pattern, such that there were 12 sample replicates per donor per plate. Positionally, each replicate had a different location within the plate layout to minimize biasing the model with plate effects, and the outer two rows and columns on the plate were only seeded with media to avoid edge effects. In total, the Hamilton system was able to create five replicate 384-well imaging plates (CellVis, P384-1.5H-N). Cells were stored in an automated incubator (Cytomat, Thermofisher) before being treated with various compounds, stained, and imaged, as described below.

### 5.3 Drug dispensation

Based on prior work and previously published data^17^, three drugs were determined to be of interest for the high-throughput screen due to prior success in shifting the cellular morphology of diseased primary human fibroblasts: Y-39983 HCl (CAS: 173897-44-4), Pemigatinib (INCB054828) (CAS: 1513857-77-6), and CP21R7 (CP21) (CAS: 125314-13-8). Stocks of the compounds of interest (10mM) were thawed and diluted to intermediary stocks of 1 mM and 0.2 mM in DMSO (D2650, Sigma). An I.Dot 2.0 Dispenser (Cellink, SO20-7799) was used to dispense 50 nL of either 1 mM or 0.2 mM stock across all five plates (final concentrations of 1 µM or 0.2 µM respectively). The final drug layout across the plates was determined algorithmically to ensure an unbiased approach with an equivalent number of replicates and well volume per experimental condition. The I.Dot additionally dispensed an equivalent volume of DMSO on top of the wells that did not receive any compound to ensure that all wells (control or compound treated) contained a total of 0.1% DMSO. Once the wells were treated with either compound or DMSO control, they were placed back into a Cytomat where they were imaged overnight on the Opera Phenix™ High Content Screening System. The plates were allowed to incubate for 72 hours, before they were removed for fixation and staining, as described below.

### 5.4 Experimental layout

The seeding system (described above) was designed to stamp a pattern 4 times from a 96 well plate to a 384. Two columns and rows were cut out from the outer rim edge to avoid edge effects leaving the plate with 12 well replicates per cell line. Each stamp was replicated 4 times, leaving a layout with 4 adjacent wells with the same cell line. The drugs were then evenly distributed among the cell lines, but for each of the 4 neighboring wells with the same cell line there was at least 1 DMSO, to balance local position effects. We generated a total of 5 plates, with 300 wells seeded with DMSO and 150 wells per condition.

### 5.5 Staining and imaging

We were then able to harness high-content image-based assays for unbiased morphological profiling with multiplexed fluorescent dyes. The original cell painting protocol^15^ was adapted to fluorescently label the cells using an automated liquid handling system, as previously described^7^. The plates were placed onto the deck of the Hamilton STAR Liquid Handler. Spent media was aspirated and replaced with FEM containing MitoTracker (Invitrogen™ M22426). Plates were then incubated at 37 °C for 30 minutes, followed by fixation in 4% Paraformaldehyde (Electron Microscopy Sciences, 15710-S) before being washed with 1x HBSS (Thermo Fisher Scientific, 14025126). The plates were then permeabilized in 0.1% Triton X-100 (Sigma-Aldrich, T8787) diluted in 1x HBSS (Thermo Fisher Scientific, 14025126). Cells were stained at room temperature with the Cell Painting staining cocktail for 30 min after two additional washes in 1x HBSS. The stain cocktail included Hoechst 33342 trihydrochloride, trihydrate (Invitrogen™ H3570), Molecular Probes Wheat Germ Agglutinin, Concanavalin A, Alexa Fluor® 488 Conjugate (Invitrogen™ C11252), SYTO® 14 Green Fluorescent Nucleic Acid Stain (Invitrogen™ S7576), Alexa Fluor® 568 Phalloidin (Invitrogen™ A12380), Alexa Fluor 555 Conjugate (Invitrogen™ W32464). Plates were washed three times post-incubation with 1x HBSS, sealed with 1x HBSS in the wells, and refrigerated until removal for imaging.

Plates were imaged using the Opera Phenix™ High Content Screening System. The inner 240 wells of each plate were imaged using 49 non-overlapping single plane fields in non-confocal mode with a 40x objective (Water, NA 1.1). To capture the entirety of the Cell Painting panel, five channels were created with differing combinations of excitation and emission spectra for each. Specifically, 375 nm and 435-480 for Hoechst 33342, 488 nm and 500-550 for Concanavalin A488, 488 nm and 570-630 for SYTO14, 561 nm and 570-630 for WGA and Phalloidin, and 640 nm and 650-760 for MitoTracker Deep Red.

### 5.6 Benchmarking preprocessing steps

We designed a 5-fold cross-validation test-train split to avoid confounding factors and overfitting to converge to misleading readouts. The experiment was conducted using 3 different normalization methods and corrected for different confounders to select the best model. For this step, we first removed all the features that were either binary or location specific (eg. cell number, distance, etc.) to preserve their raw values before zero-averaging all of the known confounders first individually, then in combination. We next applied the three different normalization methods that we compared in the analysis: the whitening transformation^26^ – a technique where the features become uncorrelated and equivariant with the same variance equal to 1-, z-scoring the features (reducing them to 0-mean and constant standard deviation), and normalizing the data between 0 and 1.

### 5.7 Overall computation pipeline description

Image file names and paths are read into memory where relevant metadata (e.g., well, field-of-view, channel, etc) is parsed from image file paths using a generalizable, user-specified pattern. Using a specified number of random images, the background trend is computed and used for flat field correction (one matrix of values per channel). Next, the images of the channels belonging to the same tile are loaded, and if the acquisition was performed in multiple planes, the images for each channel are flattened into a max projection. The images to be processed are loaded, flattened, flat-field corrected, normalized, and converted into 8-bit grayscale. After these pre-processing steps, the segmentation step is performed to locate the cells and start the single-cell analysis. This step can be performed using multiple different approaches. For this paper, as the dataset we used consisted of fibroblasts, which are very easy to separate, we used a Triangle threshold algorithm^39^ on the nuclei channel. For each of the detected cells, a crop of a specified size (in this case 598×598) was segmented from the centroid of the nuclei, in order to minimize computational time and ensure consistency in our measurements across cells. By doing so, we were able to maintain a uniform reference for all the cells, resulting in more comparable and homogeneous measurements.

For each of the channels, there is first a further segmentation step to identify the mask of the tagged portion of the cell within the crop, then channel-specific measurements are performed. In cases where multiple cells would be within the same crop, only the mask overlapping with the nuclear mask was considered part of the cell, and for the DNA channel, we used the mask that included the center pixel of the crop. The output of the pipeline is a csv file for each plate containing a row for each cell’s computed feature vector. The number of files can be changed if the user specifies a maximum file size. More details about the code and the specific usage can be found in the publicly available repository (https://github.com/NYSCF/ScaleFEx).

#### 5.7.1 Features definition

For each channel, a set of different measurements is extracted starting from the initial segmentation (Figure 1, Supplementary Figure 1a). The main categories of measurements are shape, texture, granularity, intensity, concentric measurements, overlap between channels, optional measurements specific to the mitochondria and RNA, and global annotations. All of the measurements are summarized and described in Supplementary Table 1. The total number of features is 1861 for a cell painting panel (5 channels), but it can vary depending on the number of dyes used and if the extra features for Mitochondria or RNA are selected. The distribution of summed features by category are shown in Figure 1 (Note that some features belong to more than 1 category, so they’re counted twice).

#### 5.7.2 Shape

Shape measurements are the features related to the shape and size of the cell. They are computed based on the per-channel segmentation mask, and measure features such as area, perimeter, radius and compactness. Shape measurements are important in understanding the high-level differences between populations and are the easiest to validate by eye.

#### 5.7.3 Texture

Texture is a measure of a cell’s definition, alongside the relative measure of dye taken up by any given cell. Texture is measured by computing a gray level co-occurrence matrix (GLCM), represented as a histogram describing the spatial relationship between pixels in an image and their gray levels. The GLCM is calculated by counting the number of times pairs of pixels with specific gray levels occur in a defined spatial relationship (calculated over multiple angles and distances). The resulting matrix is used to extract features that describe the texture, directionality, and other properties of the image. We computed texture for 5 distances and for 5 different angles. To avoid background biases, we computed these measurements on the image multiplied by the mask (the background results in 0 values).

#### 5.7.4 Granularity

Granularity in an image refers to the level of detail and resolution of its components. It is a measure of the size of the individual elements that make up an image, and how well these elements are defined and separated. A high-granularity image has a fine resolution with clearly defined elements, while a low-granularity image has a coarse resolution, and its elements appear blurred or blended. To measure granularity, a deconvolution process can be performed using round kernels of varying sizes on the image, and the average value can be calculated. Similar to the texture measurements, we assigned a 0 value to all the pixels outside the segmentation mask.

#### 5.7.5 Intensity

Intensity measurements refer to the measurement of the brightness or darkness of individual pixels in an image. The intensity of a pixel is represented by a numerical value that corresponds to the amount of light that the pixel reflects or absorbs. The intensity measures help to identify if some dyes are particularly reactive or condensed on some cells with respect to others.

#### 5.7.6 Concentric measurements

Concentric measurements are intensity and granularity measurements computed in a radial fashion. Concentric areas are used to define new masks to overlay to the image and compute over the reduced images the measurements described above. This set of features help to identify differences in the radial distribution within the cells.

#### 5.7.7 Zernike moments

Zernike moments utilize a set of complex polynomials that are orthogonal over the unit disk, providing a robust representation of image features. The computation of these moments was facilitated by the mahotas library, which provides an efficient implementation for their calculation.

#### 5.7.8 Overlap between channels

This set of measurements calculates the relationships between different channels and how much they correlate to each other. They help highlight differences in the cell’s compartments.

#### 5.7.9 Mitochondria

This optional measurement is specific to the mitochondria channel. In computing this set of measurements, an additional segmentation step is performed within the mask, followed by a skeletonization to highlight the fine net-structures of mitochondria. From this second binary mask, volumetric information is extracted, and from its skeletonization (Supplementary Figure 1a), we can calculate the total extent of mitochondria, the number of branches and the number of endpoints.

#### 5.7.10 RNA

Similar to the mitochondria measurements, the RNA measurements are also optional and calculate the number and size of the nucleoli overlaid to the DNA mask. This measurement is performed through an extra step of segmentation (Supplementary Figure 1a) and includes counts of organelles and total volume.

#### 5.7.11 Global

Global measurements include the information retrieved at tile level and are the same for all the channels. They log the density of the site (number of cells), and distance from the 2 closest cells. Having knowledge about the spatial surroundings of the cells is very important, as the cells can assume very different phenotypes depending on density and proximity to other cells.

### 5.8 Code description

The ScaleFEx℠ algorithm and its associated modules are publicly available on the NYSCF GitHub repository (https://github.com/NYSCF/ScaleFEx), which also includes comprehensive documentation in its Wiki section. This repository hosts the code and detailed instructions for replicating the analyses.

The algorithm processes high-content imaging (HCI) screens to extract a fixed vector of features. The system supports deployment both on-premise and on AWS cloud infrastructure, catering to various computational needs.

Initialization and Configuration: A Python class named “Process_HighContentImaging_screen” (“Process_HighContentImaging_screen_on_AWS for the AWS version) initializes with parameters from a YAML configuration file specifying directories, experiment details, and processing parameters. The class leverages multiprocessing for efficient data processing. Features are stored in a pandas dataframe for subsequent analysis. To make sure that the computation was successful, we record a file that for each well and site logs the number if cells that were computed, and the ones that were not together with the reason (eg. the cell was near the border, or the segmentation failed). This file is also used to retrieve the computation from the last locations, instead of starting from scratch again.

AWS Integration: To facilitate the deployment of ScaleFEx℠, two AWS CloudFormation stacks were created. The first stack sets up essential security infrastructure, including a Virtual Private Cloud (VPC), subnets, and a security group, along with the necessary IAM permissions. These components are referenced by the subsequent template, AWS_CloudFormation_Templates/ScaleFEx_init.yaml. This second stack initializes a master EC2 machine (c5.12xlarge) and executes the AWS_scalefex_main.py python script. This machine orchestrates the workflow leveraging AWS’ SDK, boto3. It launches and monitors multiple spot worker machines which run the ScaleFEx_extraction.py script on different parts of the dataset in parallel. Throughout this process, images are processed and the resulting data is securely stored and managed within an AWS S3 bucket.

Data Query and Preprocessing: The system queries image data, applies flat field corrections, and normalizes images. It is equipped to handle images of specified sizes and adjusts for the maximum and minimum cell sizes. The data is queried using multiple parameters that detail the image directory structure, specific sets of images to be analyzed (e.g., plates, measurements, etc.) and how relevant metadata is encoded within the directory and file names. Specifically, we devised a pattern-based querying function which takes a string input detailing the order, character length, and names of metadata fields that is generalizable to virtually any imaging dataset.

Feature Extraction: The ScaleFEx_from_crop module includes all the functions used for computing the features starting from a crop image of a cell, including mask computation, shape feature extraction from segmented regions, texture analysis, granularity, intensity measurements, zernike moments, concentric measurements and correlation across multiple channels.

### 5.9 Analysis details

#### 5.9.1 In-house dataset analysis

A Jupyter notebook is provided in the GitHub repository together with the main code. The output of ScaleFEx℠ is a csv file (or a parquet file if the AWS version was used) for each plate containing 1 cell’s feature vector per row, along with location information (e.g., well, field, coordinates). After using the pandas library to load the csv or parquet, for the final analysis we normalized all the values between 0 and 1 and averaged at well level. We next used a LR algorithm (sklearn.linear_model.LogisticRegression) to predict drug vs control and evaluated the result for each confounder removed. The model was designed in 5 Cross Validation (CV) folds with a plate held out for testing for each fold, on data normalized between 0 and 1. To remove the contribution of any given confounder, we removed the mean of each one and re-normalized at the end of the process to keep the values between 1 and 0, as previously described^7^. We visually assessed the data using a UMAP (from umap library). Variance thresholding was performed (sklearn.feature_selection.VarianceThresholding) to remove any 0-variance features that might occur. To determine how many features were necessary to score the best performance, we used a LR model and max iteration of 100000 adding 1 feature at a time for features sorted by importance to select until the model reached its peak accuracy. Once the number of features was found for every drug, we used a Recursive Feature Elimination model (sklearn.feature_selection.RFE) with a LR estimator to select the best necessary features. From a LR algorithm, we ranked the features (model.coeff_), removed the highly correlated ones feeding them in the ranked order filtering by Peorson’s correlation coefficient 0.9 >, and checked how many times the same feature occurred within the cross validation folds. The result is summarized in Supplementary Table 2. Seaborn and matplotlib libraries were used to visualize the heatmap and the data distributions.We employed a two-sided Mann–Whitney U-test to assess the statistical differences between two classes. To account for multiple comparisons, we applied a Bonferroni adjustment to the significance levels. This method was chosen for its nonparametric nature, which does not assume a normal distribution of the data, making it suitable for our analysis where the distribution of data could not be assumed to be normal. The images of cells were extracted by selecting the well with the feature of interest closest to its mean. A 3×3 tile portion of the well was visualized to maintain a decent resolution.

#### 5.9.2 RxRx2 dataset analysis

The analysis was performed using a similarity matrix derived from the data loaded into a pandas dataframe, where each element represents the similarity between each well vs each other. The control samples (marked as “EMPTY”) were used as baselines for comparing the effects of each small molecule. Conditions showing significant deviations were identified by comparing their mean similarity scores against the mean plus standard deviation of the control group’s similarity scores. Specifically, conditions were flagged if their mean similarity was less than the mean of the control group minus one standard deviation of the control group. In a similar way, the compounds were grouped based on the similarity between each other. The meaningful features were then extracted as described in the paragraph above.

## 6. Acknowledgements

This work was supported by The New York Stem Cell Foundation (NYSCF). We thank members of the NYSCF Global Stem Cell Array® Team for cell expansion handling and method development that led to the final experiment. We are grateful to Sandra Capellera-Garcia, Corvis Richardson and Raeka Aiyar for manuscript input and review. We thank Bjarki Johannesson for the discussions and contribution at the start of the project. We are grateful to Stefan Semrau for helping with text revision and technical input. We are grateful to all the study participants who donated samples for this research. This work was made possible through support from the Michael J Fox Foundation (MJFF-021530), The Silverstein Foundation, a National Academy of Medicine’s Healthy Longevity Catalyst Award, and NYSCF.

## 7. Code and Data Availability

All code will be made available under the BSD-3-Clause Clear license and shared through this GitHub repository: https://github.com/NYSCF/ScaleFEx. Raw imaging data is hosted on an Azure bucket and can be downloaded using this link: https://nyscfopensource.blob.core.windows.net/nyscfopensource/scalefex/ScaleFExDataset.zip. CSV files of ScaleFEx raw features already computed on this dataset can be downloaded here: https://nyscfopensource.blob.core.windows.net/nyscfopensource/scalefex/scalefex_raw_features.p <u>arquet</u>

The corrected, normalized and with uncorrelated features well averages can be found in the above mentioned GitHub repository.

## 9. Figure captions

Supplementary Figure 1

a) Example of segmentation masks for each channel in a single cell and additional segmentation masks used for RNA and Mitochondria (skeleton only) and illustration to describe the additional features related to Mitochondria and RNA that are computed. An additional mask was first computed to segment the sub-structures of the channel. For RNA, the values related to the number of Nucleoli and volume were calculated, shown in the mask (bottom right, panel b – RNA Mask).

For mitochondria, measurements related to volume on the mask (purple portion in the schematics) are first computed, before computing the mask’s skeleton and subsequently branches, endpoints, total length and number.

b) Experimental pipeline: Using a collection of skin fibroblasts from a cohort of 20 donors, 5, reproducibly seeded, 384 well plates were generated for compound profiling. Three drugs at 2 different concentrations were applied, alongside DMSO controls via automated dispensing. After 72 hours of growth in the presence of the drugs, the cells were fixed, Cell-Painted and imaged using an automated microscope.

Supplementary Figure 2

UMAP of the feature vectors color-coded by drug and condition. Each column shows a different normalization method, and each row is for each different model.

Supplementary Figure 3

Value distribution and image visualization of the most important features to classify Pemigatinib at a) 0.2 μM and b) 1.0 μM. The pictures depict the wells with the closest value to the mean of the feature. In interest of space, we only visualize 9 out of the 49 total tiles that make a well. In the heatmaps, the darker the color, the higher the number of features. The number states the actual number of features per class and channel. The half numbers are for those features with double class or double channel. The feature distribution values are all normalized between 0 and 1. Boxes represent median, upper and lower quartiles, the whiskers represent the standard deviation, and the diamonds the outliers. **** = p < 0.0001.

Supplementary Figure 4

Value distribution and image visualization of the most important features to correctly classify CP21R7 at a) 0.2 μM and b) 1.0 μM. The features of interest are shown on top of the images. The pictures depict the wells with the closest value to the mean of the feature. In interest of space, we only visualize 9 out of the 49 total tiles that make a well. In the heatmaps, the darker the color, the higher the number of features. The number states the actual number of features per class and channel. The half numbers are for those features with double class or double channel. The feature distribution values are all normalized between 0 and 1. Boxes represent median, upper and lower quartiles, the whiskers represent the standard deviation, and the diamonds the outliers. **** = p < 0.0001.

Supplementary Figure 5

Value distribution (box plots) and image visualization of the most important features that distinguish similar drug groups from the control (Group 1,2 and 3), and groups with unique effects (Group 4). The features of interest are shown on the x axis of the box-plot. The pictures depict the wells with the closest value to the mean of the feature. In interest of space, we only visualize 9 out of the 49 total tiles that make a well. The feature distribution values are all normalized between 0 and 1. Boxes represent median, upper and lower quartiles, the whiskers represent the standard deviation, and the diamonds the outliers. **** = p < 0.0001.

## 10. Table captions

Supplementary Table 1

Summary of all the features computed by ScaleFEx℠. Each column specifies the feature name, category, the formula used to compute the feature, the package (if any) used for the computation, an explanation of their meaning and if the feature is computed by channel. We removed the channel name and the rounds of features (but one) that went through multiple iterations to avoid redundant information in the table.

Supplementary Table 2

Accuracy and AUC of the binary classifiers built to identify treated VS control wells. The values are computed using models based on ScaleFEx℠, CellProfiler and embeddings and across all the different combinations of confounder removal and normalization.

Supplementary Table 3

Summary of the best scoring features for each drug. Each sheet contains the features relative to the drug and concentration specified in the sheet-name. The features are sorted by the total counts of features making the cut for each plate and the sum of the coefficients. In the remaining columns there are the coefficients for each of the plates

Supplementary Table 4

Tab1: List of analyzed small compounds from the RxRx2 dataset

Tab2: Groups of the drugs with similar effects and statistically different from the control

