## supplementary_figures for "A Highly-Efficient, Scalable Pipeline for Fixed Feature Extraction from Large-Scale High-Content Imaging Screens"

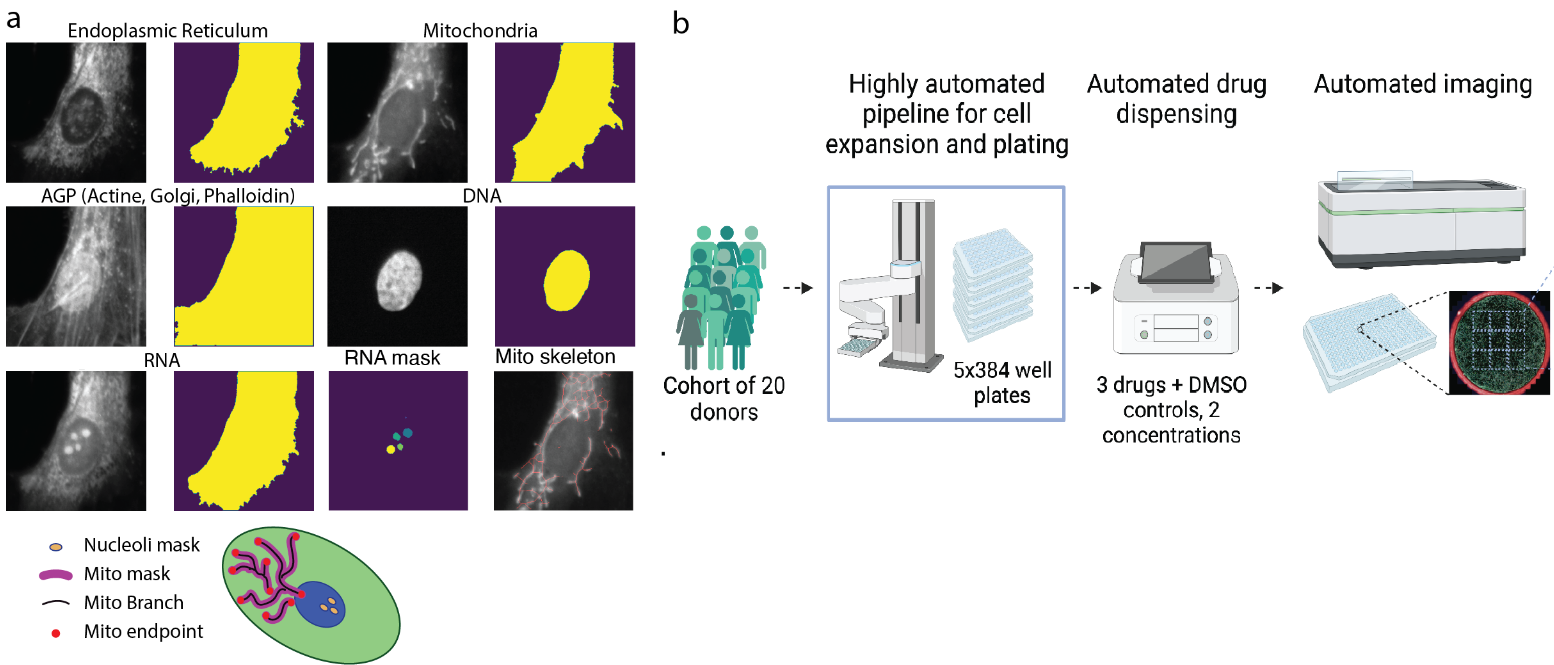
a
Endoplasmic Reticulum
Mitochondria
AGP (Actine, Golgi, Phalloidin)
DNA
RNA
RNA mask
Mito skeleton
Nucleoli mask
Mito mask
Mito Branch
Mito endpoint
b
Highly automated pipeline for cell expansion and plating
Automated drug dispensing
Automated imaging
Cohort of 20 donors
5x384 well plates
3 drugs + DMSO controls, 2 concentrations

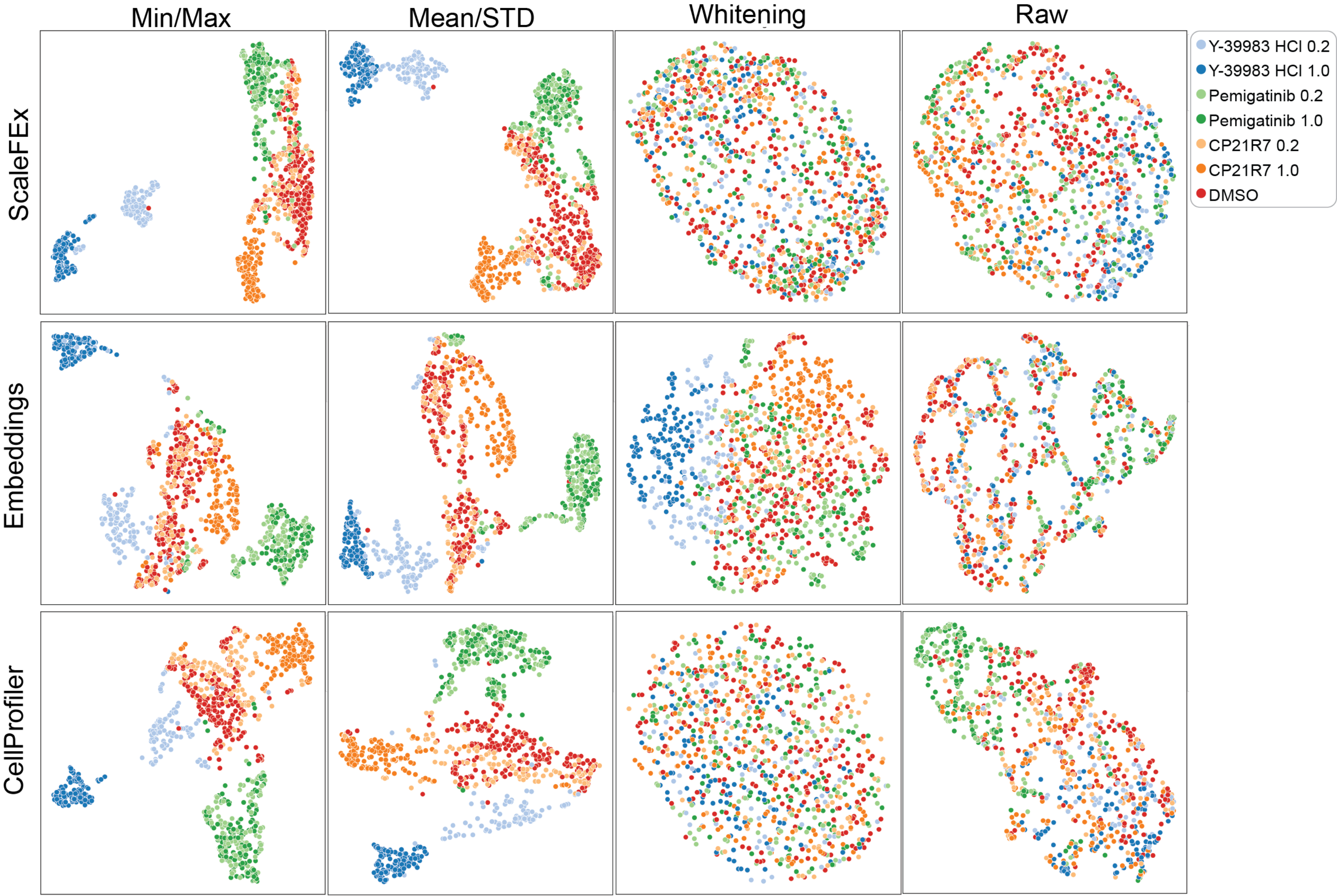
Min/Max
Mean/STD
Whitening
Raw
ScaleFEx
Embeddings
CellProfiler
Y-39983 HCl 0.2
Y-39983 HCl 1.0
Pemigatinib 0.2
Pemigatinib 1.0
CP21R7 0.2
CP21R7 1.0
DMSO

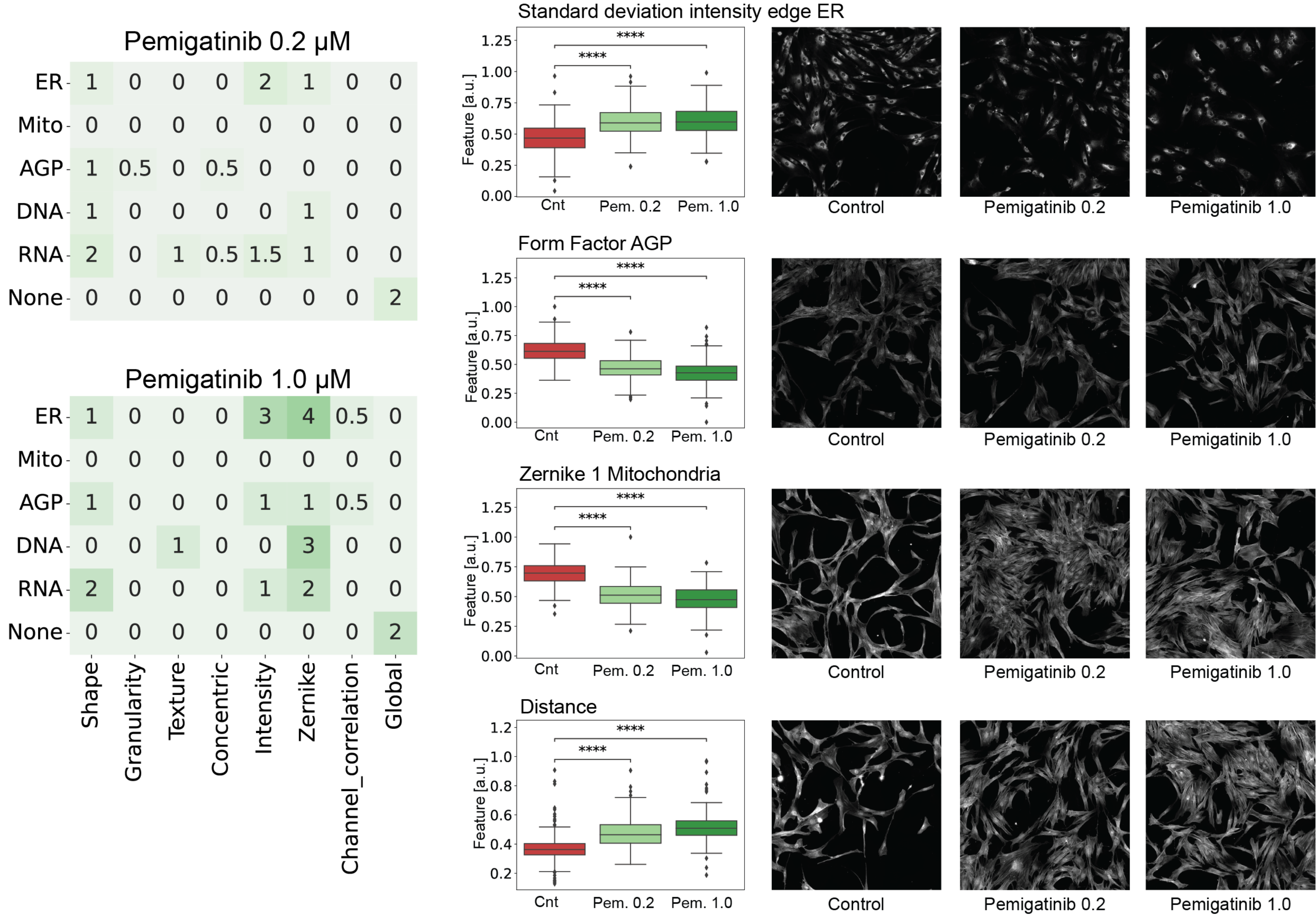

Pemigatinib 0.2 µM
ER 1 0 0 0 2 1 0 0
Mito 0 0 0 0 0 0 0 0
AGP 1 0.5 0 0.5 0 0 0 0
DNA 1 0 0 0 0 1 0 0
RNA 2 0 1 0.5 1.5 1 0 0
None 0 0 0 0 0 0 0 2
Pemigatinib 1.0 µM
ER 1 0 0 0 3 4 0.5 0
Mito 0 0 0 0 0 0 0 0
AGP 1 0 0 0 1 1 0.5 0
DNA 0 0 1 0 0 3 0 0
RNA 2 0 0 0 1 2 0 0
None 0 0 0 0 0 0 0 2
Shape
Granularity
Texture
Concentric
Intensity
Zernike
Channel_correlation
Global
Standard deviation intensity edge ER
Form Factor AGP
Zernike 1 Mitochondria
Distance
Feature [a.u.]
****
Cnt
Pem. 0.2
Pem. 1.0
Control
Pemigatinib 0.2
Pemigatinib 1.0

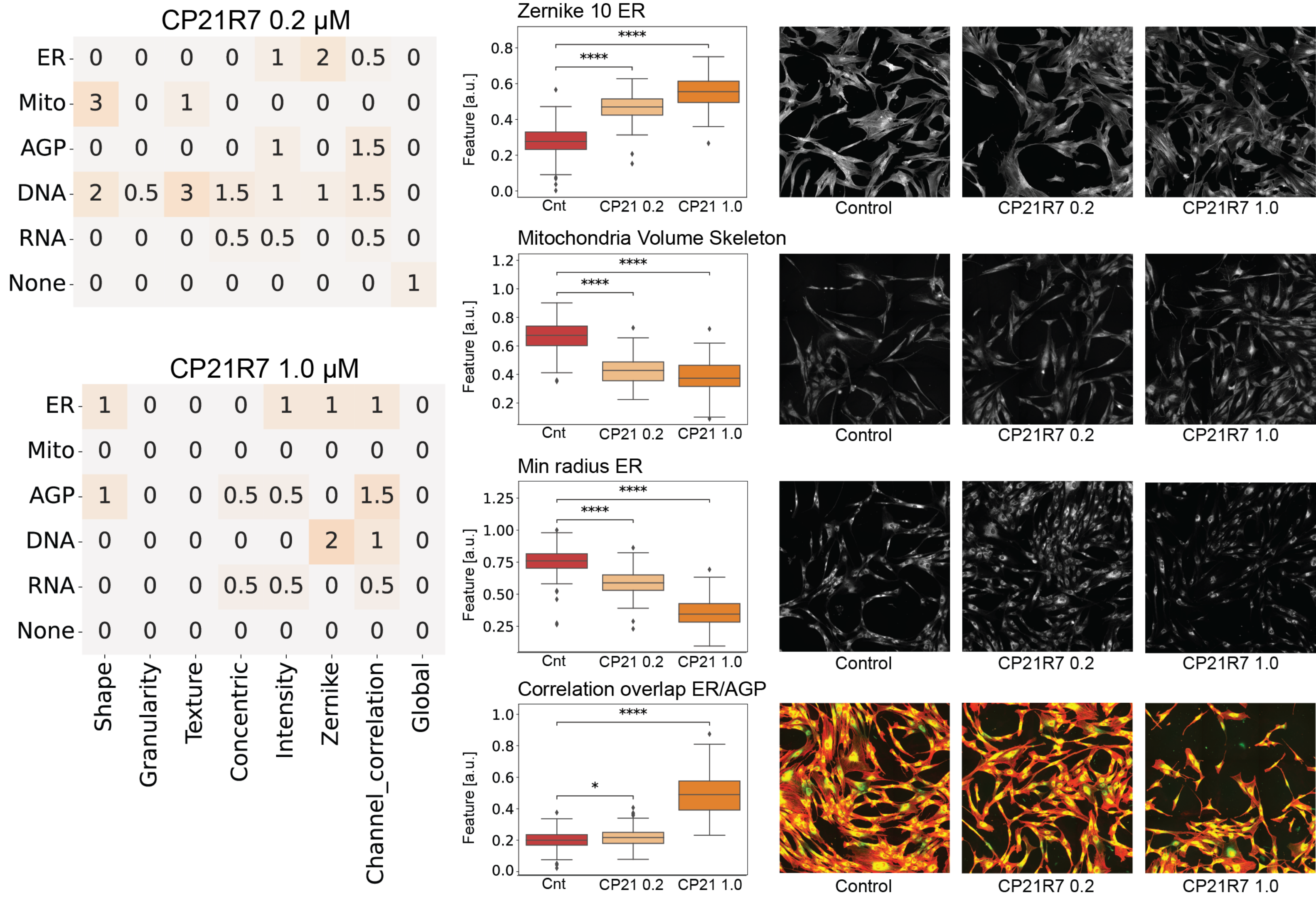
CP21R7 0.2 μM
ER 0 0 0 0 1 2 0.5 0
Mito 3 0 1 0 0 0 0 0
AGP 0 0 0 0 1 0 1.5 0
DNA 2 0.5 3 1.5 1 1 1.5 0
RNA 0 0 0 0.5 0.5 0 0.5 0
None 0 0 0 0 0 0 0 1
CP21R7 1.0 μM
ER 1 0 0 0 1 1 1 0
Mito 0 0 0 0 0 0 0 0
AGP 1 0 0 0.5 0.5 0 1.5 0
DNA 0 0 0 0 0 2 1 0
RNA 0 0 0 0.5 0.5 0 0.5 0
None 0 0 0 0 0 0 0 0
Shape
Granularity
Texture
Concentric
Intensity
Zernike
Channel_correlation
Global
Zernike 10 ER
Mitochondria Volume Skeleton
Min radius ER
Correlation overlap ER/AGP
Feature [a.u.]
Cnt
CP21 0.2
CP21 1.0
****
*
Control
CP21R7 0.2
CP21R7 1.0

### Group 1: drugs with similar effects

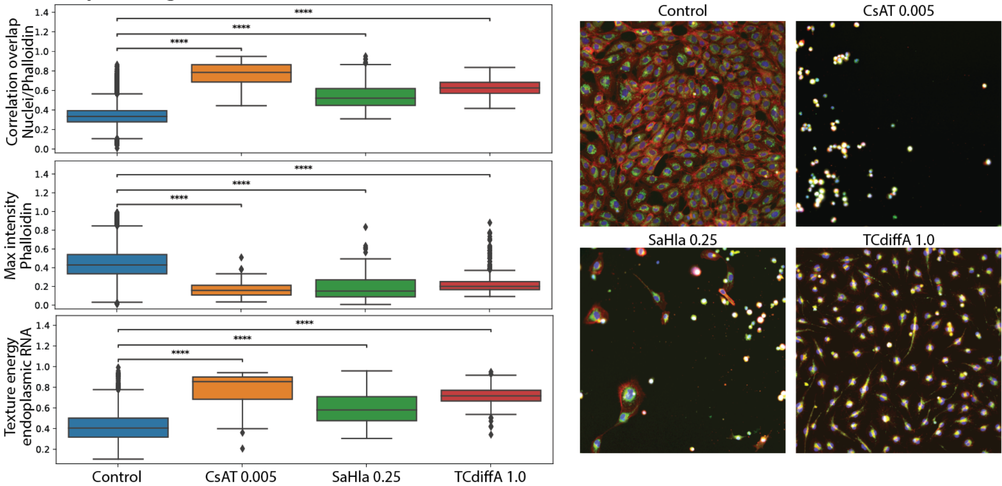


### Group 2: drugs with similar effects

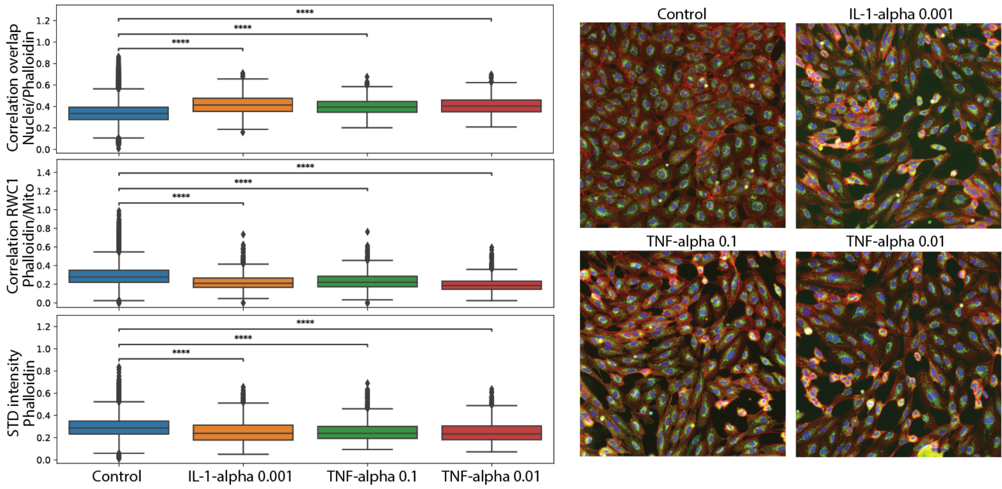


### Group 3: drugs with similar effects

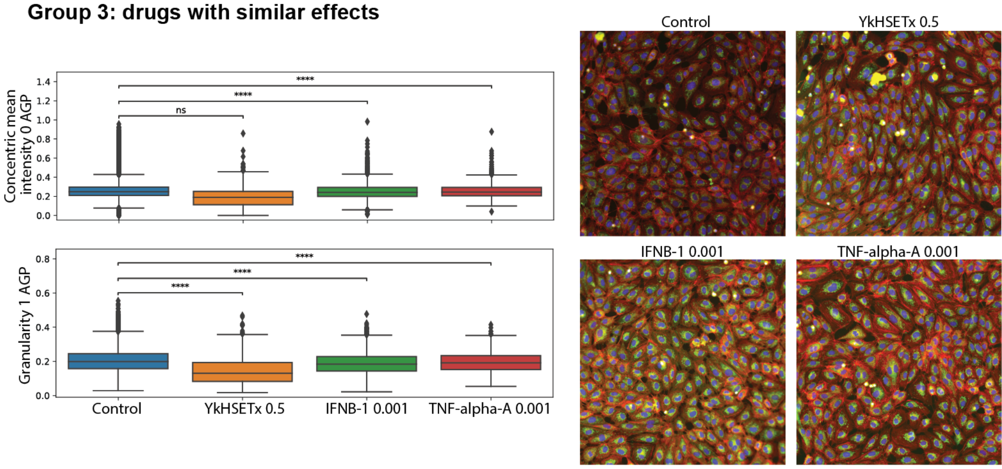


### Group 4: example of drugs with unique effects

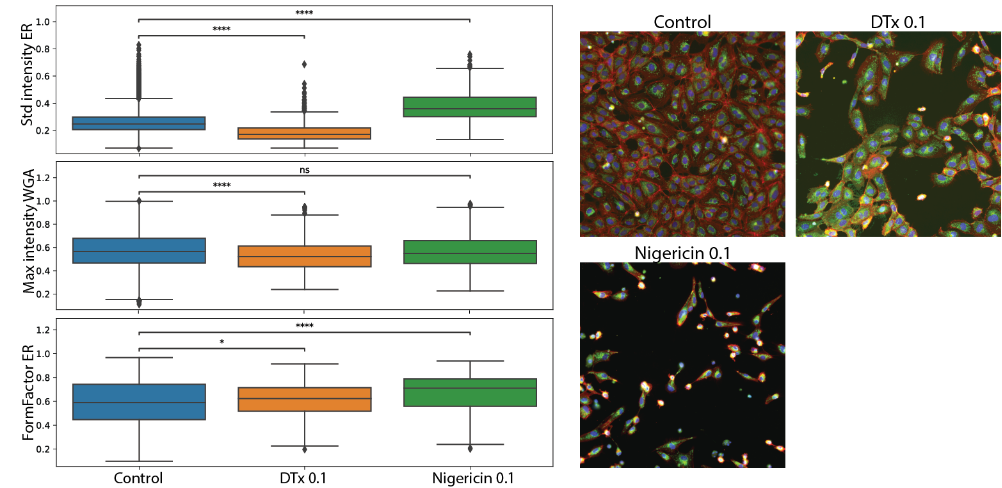
